# Elevated temperature alters microbial communities, but not decomposition rates, during three years of in-situ peat decomposition

**DOI:** 10.1101/2023.04.13.536719

**Authors:** Spencer Roth, Natalie A. Griffiths, Randall K. Kolka, Keith C. Oleheiser, Alyssa A. Carrell, Dawn M. Klingeman, Angela Seibert, Jeffrey P. Chanton, Paul J. Hanson, Christopher W. Schadt

**Affiliations:** Biosciences Division, Oak Ridge National Laboratory, Oak Ridge, TN, USA 37830; Environmental Sciences Division, Oak Ridge National Laboratory, Oak Ridge, TN, USA 37830; Climate Change Science Institute, Oak Ridge National Laboratory, Oak Ridge, TN, USA 37830; Northern Research Station, USDA Forest Service, Grand Rapids, MN, USA 55744; Department of Geosciences, Boise State University, Boise, ID, USA 83725; Department of Earth, Ocean, and Atmospheric Science, Florida State University, Tallahassee, FL, USA 32306; Department of Microbiology, University of Tennessee, Knoxville, TN, USA 37996

**Keywords:** Peatlands, climate change, microbiome, organic matter decomposition

## Abstract

Peatlands store approximately one-third of the global terrestrial carbon and are historically considered carbon sinks due to primary production outpacing microbial decomposition of organic matter. Climate change has the potential to alter the rate at which peatlands store or release carbon, and results from the Spruce and Peatland Responses Under Changing Environments (SPRUCE) experiment have shown net losses of organic matter and increased greenhouse gas production from a boreal peatland in response to whole-ecosystem warming. In this study, we utilized the SPRUCE sites to investigate how warming and elevated CO_2_ impact peat microbial communities and peat soil decomposition. We deployed peat soil decomposition ladders across warming and CO_2_ treatment enclosures for three years, after which we characterized bacterial, archaeal, and fungal communities through amplicon sequencing and measured peat mass and compositional changes across four depth increments. Microbial diversity and community composition were significantly impacted by soil depth, temperature, and CO_2_ treatment. Bacterial/archaeal α-diversity increased significantly with increasing temperature, and fungal α-diversity was significantly lower under elevated CO_2_ treatment. Trans-domain microbial networks showed higher complexity (nodes, edges, degree, betweenness centrality) of microbial communities in decomposition ladders from warmed enclosures, and the number of highly connected, hub taxa within the networks was positively correlated with temperature. Methanogenic hubs were identified in the networks constructed from the warmest enclosures, indicating increased importance of methanogenesis in response to warming. Microbial community responses were not however reflected in measures of peat soil decomposition, as warming and elevated CO_2_ had no significant short-term effects on soil mass loss or composition. Regardless of treatment, on average only 4.5% of the original soil mass was lost after three years and variation between replicates was high, potentially masking treatment effects. Many previous studies from the SPRUCE experiment have shown that warming is accelerating organic-matter decomposition and CO_2_ and CH_4_ production, and our results suggest that these changes may be driven by warming-induced shifts in microbial communities.

## **1.** Introduction

Despite covering less than 10% of the Earth’s surface, peatlands contain approximately one-third of all global terrestrial organic matter (OM) [1–3]. Peatland organic soil deposits can be several meters deep, the result of thousands of years of net primary production outpacing OM mineralization. Microorganisms are primarily responsible for OM degradation in peatlands [4, 5]; however, anoxic, acidic, oligotrophic, and cold conditions that are common across northern peatlands greatly constrain microbial activity [4].

Climate change has the potential to alter peatland biogeochemistry, especially at northern latitudes where warming is occurring at an accelerated pace compared to equatorial regions [6]. The large carbon stocks in northern peatlands that have built up over millennia may therefore be vulnerable to climate change [7]; however, the effects of warming and increased atmospheric CO_2_ on peatland ecosystems remain to be fully described [8, 9]. The Spruce and Peatland Responses Under Changing Environments (SPRUCE) experiment is a long-term warming and elevated CO_2_ experiment investigating peatland responses to climate change on an ecosystem level (https://mnspruce.ornl.gov/). Since 2016, whole-ecosystem warming up to +9 °C above ambient has been applied to a boreal peatland in a regression-based design. In addition, elevated air partial pressure of CO_2_ has been applied to half of the 10 SPRUCE experimental enclosures.

Results from the SPRUCE experiment have shown significant, rapid loss of carbon from the peatland with increasing temperature [10] concomitant with a large decline and death of *Sphagnum sp*. at the highest temperatures, that contribute the largest share of Gross Primary Production (GPP) in these ecosystems [11]. In addition, porewater concentrations of CO_2_ and CH_4_ have been shown to correlate with temperature treatment at SPRUCE [12], and radiocarbon analysis of soil suggests that warming is promoting microbial respiration of solid-phase peat [13]. Similar results have also been obtained in studies of other peatlands [14–16], and in incubation studies that show increasing CH_4_ and CO_2_ production with warming [17, 18].

Net losses of carbon from the SPRUCE sites have been attributed to increased degradation of OM, rather than a reduction in primary production [10]. Under anoxic conditions, microbial degradation of OM involves multiple steps including hydrolysis, fermentation, and anaerobic respiration. Mineralization of carbon to CO_2_ and CH_4_ in peatlands therefore relies on microbial metabolic interactions which may be altered by climate change. Incubation experiments have demonstrated that microbial activities [19, 20] and metabolic interactions are significantly altered by warming [15], although impacts on community structure and diversity vary [17, 18, 21, 22]. *In situ* results showing increasing CH_4_:CO_2_ with warming suggest that microbial interactions in the SPRUCE sites may be altered to favor increased methanogenesis [12, 23].

Previous studies have investigated the effects of warming on peat soil decomposition through incubations or whole-ecosystem assessments [10, 12, 13, 17, 18, 24]. Valuable insights have been gained from these experiments; however, incubation studies do not fully reflect environmental conditions, and “bottle effects” may influence microbial community structure [25]. Conversely, assessing decomposition from *in situ* environmental measurements is complex and may be influenced by other ecosystem processes such as changes in primary productivity. To overcome these limitations, we utilized new peat soil decomposition ladders, which are peat litter bags attached to a rigid frame, to assess the impacts of temperature, and CO_2_ treatments on peat soil decomposition at four depths. This approach allows for *in situ* investigation of decomposition while controlling for the effects of primary productivity and excluding fresh litter inputs. While studies of fresh plant litter decomposition using similar methods are quite common across many forest ecosystem types[26], studies of soil and particularly peat decomposition using decomposition bag methods appear to be absent from the literature. Peat decomposition ladders were deployed in the top 40 cm of peat in the 10 SPRUCE experimental enclosures, as changes in OM mineralization have been most pronounced in the surface and intermediate layers of peat [12, 24, 27]. Following a three-years of *in situ* incubation, we measured changes in peat soil mass and chemical composition and characterized microbial communities in the decomposition bags through amplicon sequencing and network analyses. We hypothesized that decomposition of peat soil would increase with increasing temperature, driven by changes in their microbial communities.

## **2.** Methods

### 2.1 Site description

The SPRUCE experiment is located on the S1 bog (low pH, acid organic soil environment) at the USDA Forest Service Marcell Experimental Forest, MN, USA. The SPRUCE site description, experimental design, warming, and CO_2_ treatments have been previously described in detail [28]. Briefly, above and belowground, whole-ecosystem warming has been applied in a regression-based design to 10 open-air enclosures on the S1 bog since August 2015. Enclosures are duplicated for each level of warming (+0 °C, +2.25 °C, +4.5 °C, +6.75 °C and +9 °C above ambient), and half of the enclosures receive an elevated CO_2_ atmosphere (+500 ppm).

### 2.2 Peat ladder construction, deployment, and retrieval

Organic soil used in the decomposition ladders was collected from within the S1 bog but outside of the footprint of the SPRUCE enclosures. Soil was collected from four depths using a Macaulay peat corer (0-10, 10-20, 20-30, and 30-40 cm), brought back to the laboratory, and air dried. After air drying, soil from each depth was homogenized by breaking up the dried soil using sieves, and large fragments of vegetation (i.e., roots) were removed. Soil was then weighed (2.4-4.2 g air-dry weight, corresponding to an approximate wet weight of 20 g) and placed into fine-mesh bags (7-µm mesh size) which were then heat sealed [29, 30].

The fine-mesh bags were placed into decomposition ladders which were made of acrylonitrile butadiene styrene plastic (design from J. Patrick Megonigal, personal communication, 2016). The ladders were placed vertically in the peat profile to allow for depth specific measurements of soil decomposition. Each ladder had four openings (6.5 cm wide by 9 cm tall) corresponding to the four soil depths (Figure S1). Each ladder was constructed by placing four, soil-filled, fine-mesh bags, one from each depth, between two plastic ladder holders. The ladders were lined with mesh (0.5 x 1.0 mm mesh size) to reduce abrasion of the fine-mesh bags during deployment and retrieval. The two ladder holders were closed with plastic fasteners.

The ladders were deployed on October 2, 2017 in the ten SPRUCE enclosures. Ladders were placed vertically into the peat so that the peat collected from 0-10 cm was incubated at that same depth. Three replicate ladders were deployed per enclosure and retrieval date. The replicate ladders were placed in three different hollow locations that are associated with companion litter and wood decomposition studies within an enclosure. Three replicate ladders per enclosure were deployed and immediately retrieved from the peatland for measurement of initial mass and chemistry (t=0 y; T_0_). On October 15, 2020 (t=3 y; T_f_), three replicate ladders per enclosure were retrieved to measure soil mass loss, changes in chemistry, and microbial community composition.

### 2.3 Peat soil mass loss and carbon, nitrogen, and phosphorus analyses

After retrieval, ladders were placed into individual plastic bags and were stored at 4 °C until processing (within 24 hours). Next, each fine-mesh bag was carefully removed from the ladder and the soil was weighed to determine wet mass. For the T_f_ bags, soil from each mesh bag was sub-sampled so that ∼1/2 was retained for dry mass and chemistry measurements, ∼1/4 was frozen at -20 °C for microbial community analyses, and the remainder was frozen (-20 °C) for archival purposes. Soil collected at T_0_ was not sub-subsampled as only dry mass and chemistry measurements were conducted.

Soil samples for mass loss and chemistry were air dried in a drying room (humidity <30%) for 1-2 months (until the change in mass was <5%) and then weighed. A subsample from each T_f_ sample was oven-dried to calculate an air-dry to oven-dry conversion factor and to calculate percent peat mass loss on an oven-dry basis. Peat used in construction of decomposition bags were not oven-dried prior to installation (T_0_) into the enclosures to prevent changes in organic matter quality, however, initial masses were corrected based on the air-dry to oven-dry conversion calculated from each sample at T_f_. Each air-dried sample from T_0_ and T_f_ was ground (IKA Tube Mill grinder, Wilmington, NC, USA) and a subsample was analyzed for carbon and nitrogen content using a combustion elemental analyzer (LECO-CHN628 analyzer, St. Joseph, MI, USA).

### 2.4 FTIR analysis

Fourier Transform Infrared Spectroscopy (FTIR) analysis was performed on the peat ladders to analyze the response of peat soil organic carbon fractions to enclosure treatments. The peat soil was ground into a homogenous powder using a Spex Sampleprep 5100 mixer-mill. FTIR spectra were collected using a JASCO 6800 FT-IR Spectrometer. Approximately 0.003 g of sample powder was secured onto the quartz crystal (Si/CaF2), and infrared light from wavenumbers 4000 cm^-1^ – 650 cm^-1^ was transmitted onto the sample at a resolution of 4 cm^-1^. Each spectrum was attenuated total reflection corrected and baseline corrected to account for variability in the beam penetration depth. To produce the functional data for molecular composition analysis, eight spectra per sample were averaged.

The spectra data were analyzed using Hodgkins’ normalization method [31]. Instrument and matrix variation impact on sample spectra absorbance were accounted for by dividing the baseline-corrected peak heights by the total integrated area of the spectrum. Using the maximum baseline-corrected absorbance between peak endpoints, the aromatics and carbohydrates functional group locations were identified. The normalized aromatics spectral peak heights were located at 1510 cm^-1^ and 1615 cm^-1^. The normalized carbohydrate spectral peak height was at 1040 cm^-1^. Each of these peak heights was used to calculate the percent of aromatics and carbohydrates in each sample.

### 2.5 DNA extraction and sequencing

DNA was extracted from ∼0.2 g of field-wet peat soil from all samples using the Qiagen DNeasy 96 PowerSoil Pro QIAcube HT Kit (Qiagen, Venlo, The Netherlands) following the manufacturer’s protocol. Amplicon metagenomic sequencing libraries were prepared as described in the Illumina 16S metagenomic sequencing library preparation guide (Part 15044223 Rev B) with a mixture of 515F and 806R primers for archaea/bacteria and custom primers designed to the ITS2 gene region for fungi [32, 33]. Pooled libraries for each sample type were validated on an Agilent Bioanalyzer (Agilent, Santa Clara, CA) using a DNA7500 chip, and the final library pool concentration was determined on an Invitrogen Qubit (Waltham, MA) with the broad range double stranded DNA assay. Paired end sequencing (2x 251x 8x 8) was competed on an Illumina MiSeq instrument (Illumina, San Diego, CA) using v2 chemistry. Due to low base diversity of the amplicons, PhiX control DNA was included in the sequencing run.

### 2.6 Sequence analysis

Demultiplexed 16S rRNA (V4 region) and ITS2 paired-end sequences were imported into QIIME2 (v. 2021.4) [34]. ITS2 primer sequences were removed using the *cutadapt* plugin [35], and all sequencing runs were individually denoised using the dada2 plugin [36]. Resulting feature tables and representative sequences were merged for downstream analyses. Taxonomy was assigned using the silva database (16S) [37] and UNITE database (ITS) [38]. Sequences and feature tables were filtered based on taxonomic assignments to include only Bacteria and Archaea, removing Chloroplast and Mitochondria sequences (16S) and Fungi with taxonomy assigned to the phylum level at minimum (ITS). A rooted phylogenetic tree was built for each dataset using the *align-mafft-to-fasttree* pipeline [39] in the *phylogeny* plugin.

### 2.7 Statistical analyses

All statistical analyses and figures were produced in R v. 4.1.0 with the *vegan* [40], *car*, *phyloseq* [41], *ggplot2* [42], *microbiome*, *SpeicEasi* [43], *igraph* [44], and *hilldiv* [45] packages. ITS and 16S feature tables, taxonomy, rooted phylogeny, and associated metadata were imported and analyzed as *phyloseq* objects. Based on the regression design of SPRUCE, temperature treatment (+0 °C, +2.25 °C, +4.5 °C, +6.75 °C, +9 °C) was used as a continuous variable for all statistical analyses. Soil depth and CO_2_ treatment were treated as categorical variables, and depth-specific subsets of the data were generated to investigate the effects of temperature and CO_2_ at a given depth. Samples were rarefied for α-diversity analyses (16S rarefaction depth = 9500; ITS rarefaction depth = 1000). The *hilldiv* package was used to calculate α-diversity metrics as Hill numbers (effective number of species) [46], and two-way ANOVA analysis was used to investigate the effects of temperature, soil depth, CO_2_ treatment, and the interaction between temperature and depth on α-diversity. Bray-Curtis dissimilarities [47] of total sum scaled data were calculated and used to assess the effects of temperature, depth, CO_2_ treatment, and the interaction between temperature, CO_2_, and soil depth on community composition by PERMANOVA. Principal coordinates analysis plots of Bray-Curtis dissimilarities were generated to visualize community composition. The *betadisper* function was used to assess differences in β-dispersion across depths, temperature treatments, and CO_2_ treatments.

Trans-domain networks were generated using the *SpiecEasi* and *iGraph* packages in R. Networks were constructed for each temperature treatment and included all depths within a given temperature treatment (n=24 network^-1^). Phyloseq objects were filtered by temperature treatment, and only amplicon sequence variants (ASVs) with a total sum of > 5 (16S) or > 3 (ITS) and an occurrence in > 20% of the samples were included in the network analyses. If a sample was lacking either a 16S or ITS library after filtering, the sample was excluded from analysis. Networks parameters were set as method=mb, nlambda=50, lambda.min.ratio=1e-3, and thresh=0.01. Empty nodes were removed, and networks were visualized using *phyloseq*. The number of nodes, edges, node degree and betweenness centrality were calculated for each network with the *igraph* package. Network hubs were identified by selecting nodes that had degree and betweenness centrality measures in the 90^th^ percentile, indicating high connectedness and centrality in the network. Pearson correlation was used to investigate the relationship between network topology and SPRUCE temperature treatments.

Kruskal-Wallis test was used to compare peat soil mass and composition at T_0_ to T_f_. Effect sizes for T_0_-T_f_ comparisons were calculated as (χ^2^ - 1)/(n - 2), where χ^2^ is the Kruskal-Wallis test result and n is the number of samples. Linear models were used to assess the effect of depth, temperature treatment, and CO_2_ treatment on soil mass loss and compositional changes.

### 2.8 Data availability

Amplicon sequences generated in this study have been deposited in the NCBI SRA under BioProject ID PRJNA941900. All code used in statistical analysis of the data and generation of figures is available at https://github.com/swroth/Peat_Decomp_SPRUCE.

## 3. Results

### 3.1 Community summary

Bacterial/archaeal communities were dominated by the phylum *Acidobacteriota* across soil depths and temperature treatments (28-46% mean relative abundance) (Figure S2). Near the surface (0-10 cm and 10-20 cm), *Actinobacteriota*, *Proteobacteria*, and *Planctomycetota* comprised approximately 35-45% of the bacterial/archaeal communities, while accounting for < 20% of the total community at 20-30 cm and 30-40 cm (Figure S2). Dominant phyla of the fungal communities also varied across depth, with *Basidiomycota* in highest relative abundance at 0-10 cm (∼62% average relative abundance), and *Ascomycota* dominating in the deeper three depths (∼54-68% average relative abundance) (Figure S3).

### 3.2 Microbial community composition and α-diversity are significantly influenced by temperature and CO_2_ treatments

Bacterial/Archaeal and fungal community compositions were significantly impacted by depth, temperature treatment, CO_2_ treatment, and the interactive effects of these variables (Figure 1, Table 1). Of the variables investigated, depth was the most influential factor driving compositional differences in the bacterial/archaeal and fungal communities, explaining 25% and 11% of the variation, respectively (Table 1). Temperature treatment explained 6.1% of the variation in the bacterial/archaeal community and 4.9% in the fungal community, and CO_2_ treatment accounted for 1.9% and 2.7% of the variation for both bacterial/archaeal and fungal communities respectively (Table 1). Significant interactive effects between soil depth and temperature treatment, and between temperature treatment and CO_2_ treatment were observed for the bacterial/archaeal communities (Table 1). Significant interactions between all tested variables were observed for fungal communities, such that temperature and CO_2_ treatment only influenced fungal community composition at specific depths (Table 1). No significant differences in β- dispersion were observed across soil depth, temperature treatment, or CO_2_ treatment.

**Figure 1:**
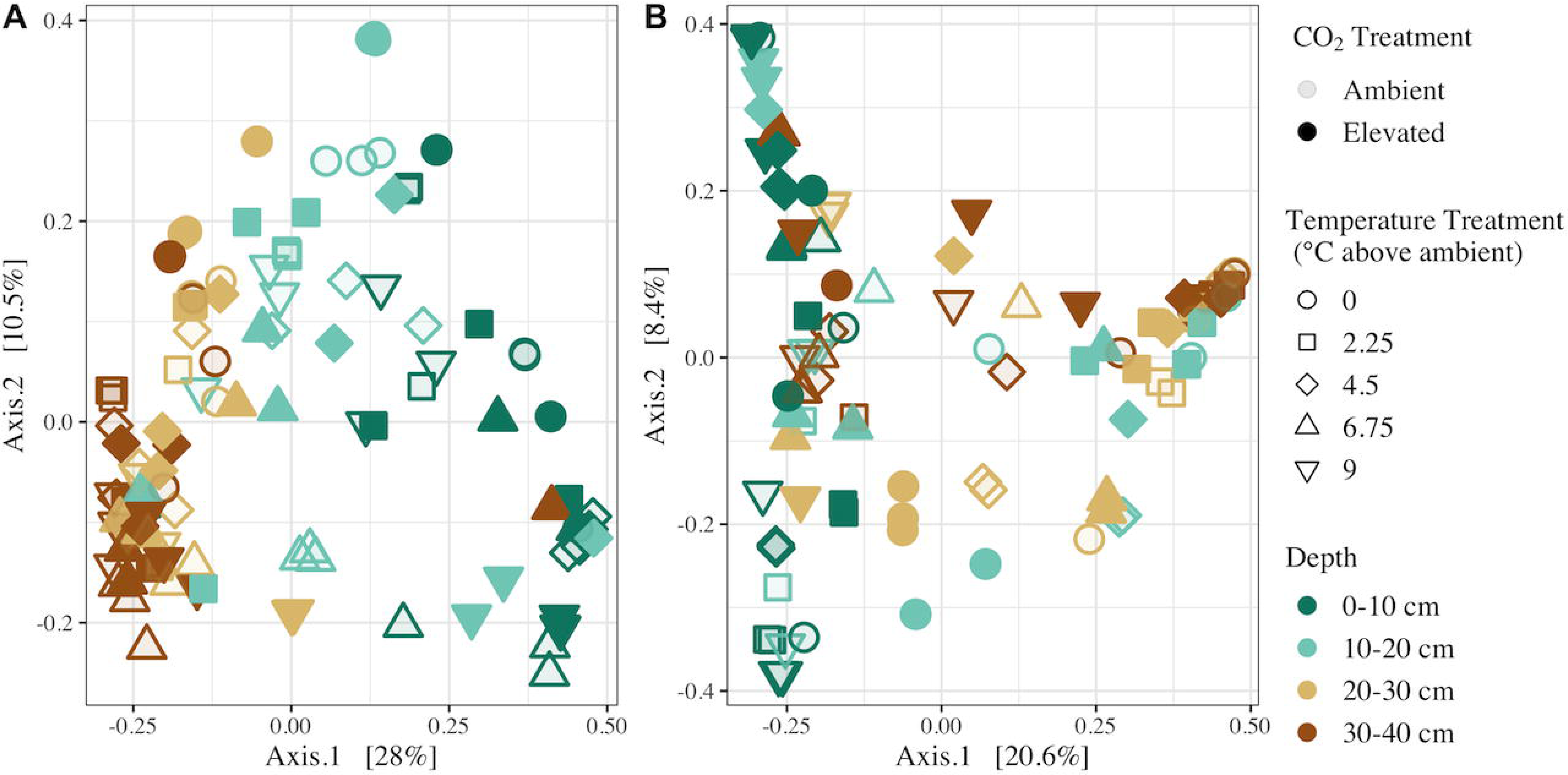
Principal coordinates analysis of Bray-Curtis dissimilarities of bacterial/archaeal (A) and fungal (B) communities in decomposition ladders. Points are colored based on sample depth, filled based on enclosure CO_2_ treatments and shaped based on enclosure temperature treatments.

**Table 1:** Results of permutational multivariate analysis of variance (PERMANOVA) of bacterial/archaeal and fungal community compositions based on Bray-Curtis distances. Significant factors and interactive effects (p < 0.05) are bolded.

| Variable | Bacterial/Archaeal |  |  | Fungal |  |  |
| --- | --- | --- | --- | --- | --- | --- |
|  | F-Statistic | R<sup>2</sup> | p-value | F-Statistic | R<sup>2</sup> | p-value |
| Depth | 15.402 | 0.25323 | **0.001** | 4.0315 | 0.10959 | **0.001** |
| Temperature Treatment | 11.1683 | 0.06121 | **0.001** | 5.46 | 0.04948 | **0.001** |
| CO<sub>2</sub> Treatment | 3.3775 | 0.01851 | **0.004** | 3.0053 | 0.02723 | **0.001** |
| Depth:Temperature Treatment | 2.907 | 0.03766 | **0.001** | 1.9049 | 0.05178 | **0.001** |
| Depth:CO<sub>2</sub> Treatment | 1.2092 | 0.01988 | 0.176 | 1.6339 | 0.04442 | **0.006** |
| Temperature treatment:CO<sub>2</sub> treatment | 4.983 | 0.02731 | **0.001** | 2.6901 | 0.02438 | **0.004** |
| Depth:Temperature Treatment:CO<sub>2</sub> Treatment | 1.4101 | 0.02318 | 0.069 | 1.8305 | 0.04976 | **0.001** |

Bacterial/archaeal α-diversity was significantly highest near the surface and declined with depth across all levels of q (species richness at q=0, Shannon entropy at q=1, Inverse Simpson at q=2) (Figure 2A, Figure S4, Table 2). Temperature treatment also significantly impacted α- diversity, with highest diversity measured in decomposition ladders from the +9 °C enclosures across all α-diversity metrics (Figure 2A, Figure S4, Table 2). Bacterial/archaeal α-diversity was significantly higher in elevated CO_2_ enclosures when compared to ambient CO_2_ at q=0 and q=1 (Figure 2A, Figure S4A, Table 2). At q=2, no effect of CO_2_ on α-diversity was observed, suggesting that CO_2_ treatment has a limited effect on abundant bacterial/archaeal taxa (Figure S4B, Table 2). In contrast, fungal α-diversity was significantly lower in enclosures with elevated CO_2_ when compared to ambient CO_2_ enclosures across all α-diversity metrics measured (Figure 2B, Figure S5, Table 2). Soil depth only significantly impacted fungal species richness (q=0), with highest richness observed at 0-10 cm, and we observed a significant effect of temperature treatment at q=2 (Figure 2B, Figure S5, Table 2) and significant interactive effects of temperature and depth at q=1 and q=2, such that temperature effects on fungal α-diversity were depth specific.

**Figure 2:**
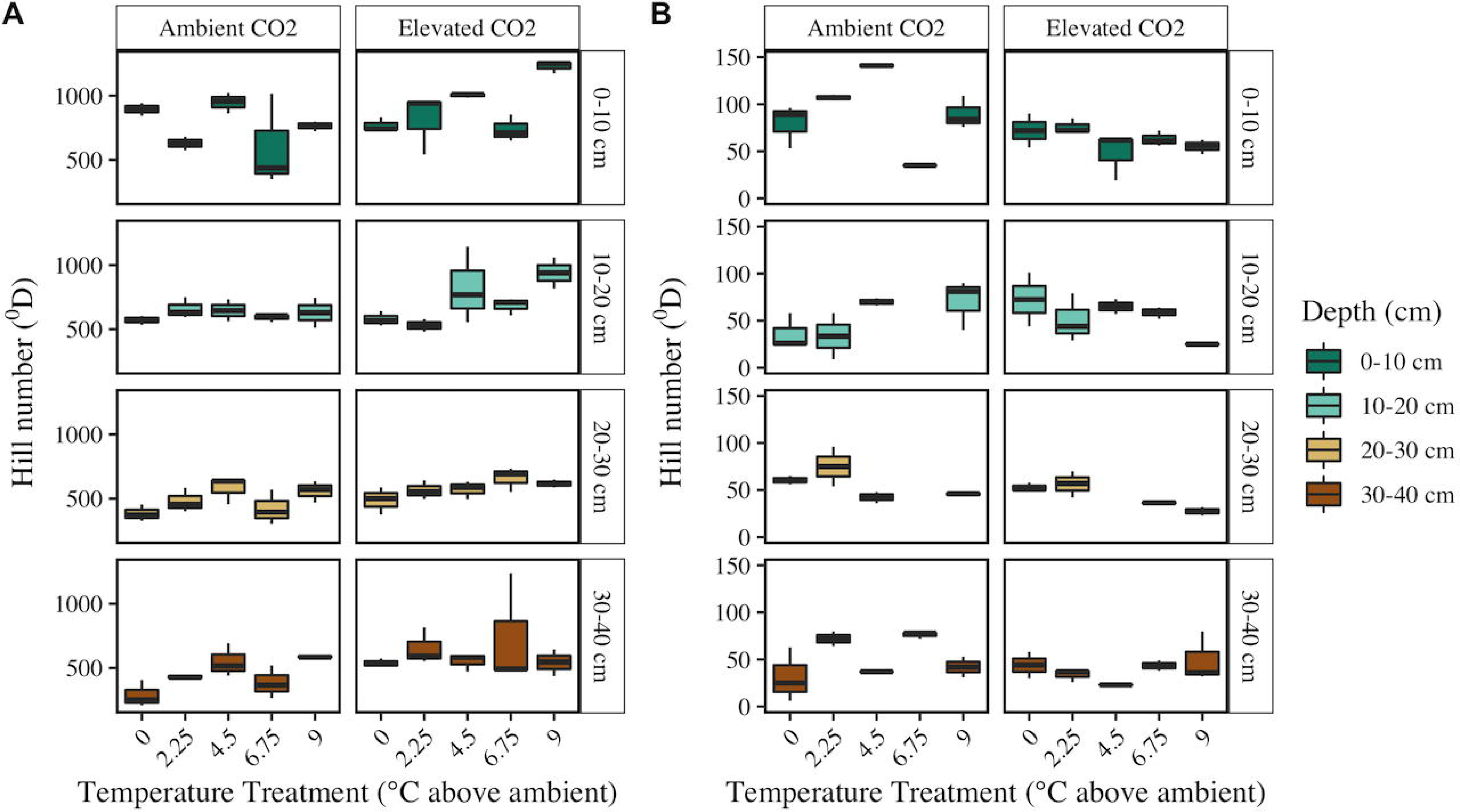
Bacterial/Archaeal (A) and Fungal (B) α-diversity at q = 0 (species richness) across depths (vertical facets/colors), CO_2_ treatments (horizontal facets), and temperature treatments (x- axis).

**Table 2:** Resulting p-values from ANOVA describing the effects of depth and SPRUCE treatments on bacterial/archaeal and fungal α-diversity across Hill numbers (q=0, q=1, and q=2). Significant values (p < 0.05) are bolded.

| Variable | 16S |  |  | ITS |  |  |
| --- | --- | --- | --- | --- | --- | --- |
|  | q=0 | q=1 | q=2 | q=0 | q=1 | q=2 |
| Depth | **<0.001** | **<0.001** | **<0.001** | **<0.001** | 0.802 | 0.296 |
| Temperature Treatment | **0.002** | **<0.001** | **<0.001** | 0.728 | 0.113 | **0.034** |
| CO<sub>2</sub> Treatment | **<0.001** | **0.015** | 0.133 | **0.009** | **<0.001** | **0.002** |
| Depth:Temperature Treatment | 0.978 | 0.907 | 0.856 | 0.094 | **0.013** | **0.003** |

### 3.3 Trans-Domain networks

We constructed temperature treatment-specific, trans-domain networks to investigate the impact of temperature treatment on the connectedness of microbial communities through comparisons of network topology (Figure S6). The microbial network from +0 °C enclosures across all depths had the lowest number of nodes (taxa), edges (connections between taxa), and average degree (mean number of edges per node), collectively indicating lower complexity of the microbial communities in +0 °C enclosures when compared to warmed enclosures (Figure 3). The number of nodes and edges and the average degree peaked in the network built from +4.5 °C enclosures (Figure 3A,B,C). Networks built from all the temperature treatments included bacterial, archaeal, and fungal nodes, with the highest number of archaeal nodes present in the +9 °C network (35 nodes), and the most fungal nodes occurring in the +4.5 °C network (126 nodes) (Figure 3B).

**Figure 3:**
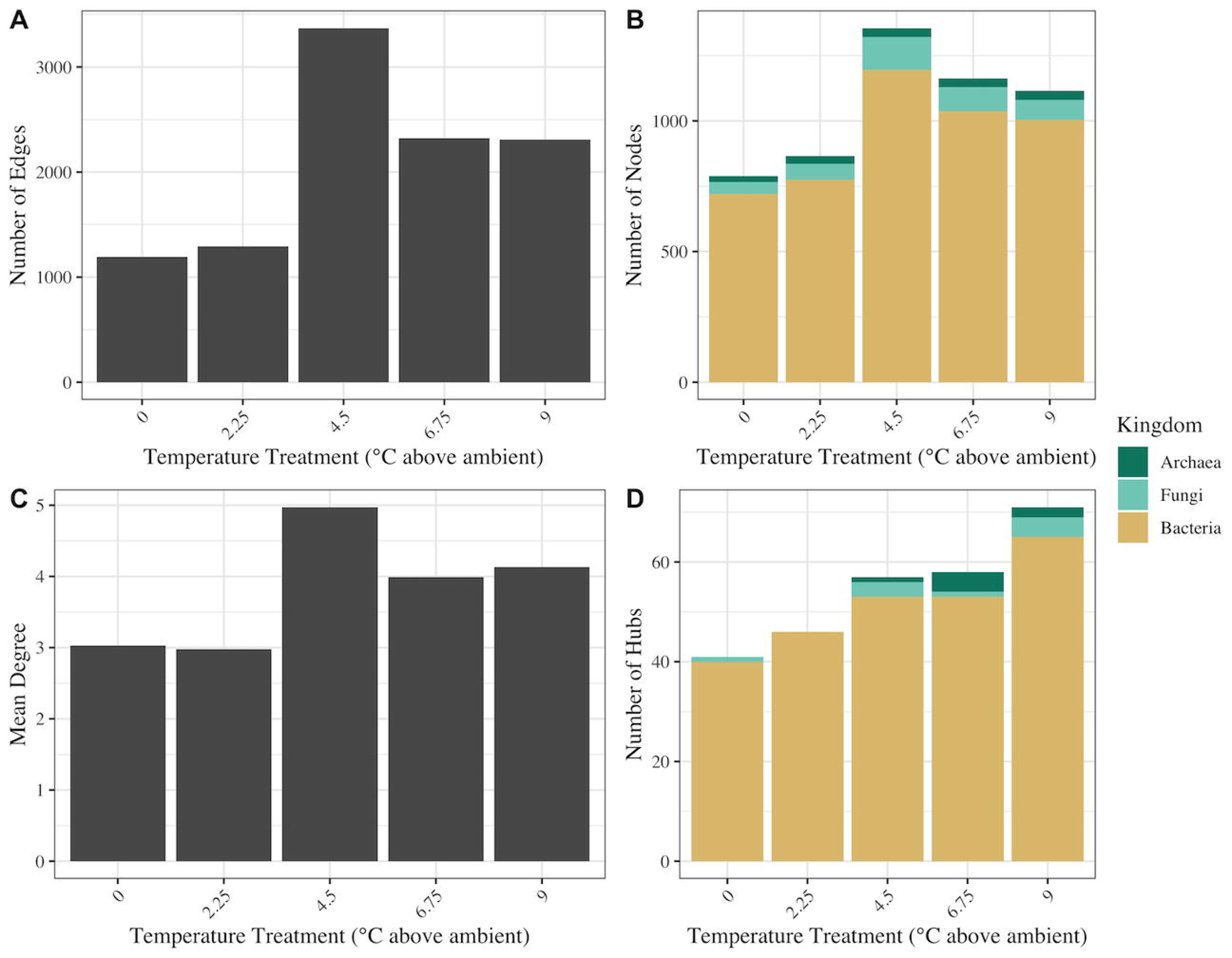
Summary of network topologies for temperature treatment microbial networks. The number of edges (A) represents the sum total connections between taxa (nodes; B) within the network. Mean degree (C) represents the average number of connections per taxon. Hubs (D) are defined as nodes within the network that are in the 90^th^ percentile for degree and betweenness centrality and represent highly connected and centralized taxa.

Within each temperature treatment network, we identified hub taxa as those with degree and betweenness centrality measures in the 90^th^ percentile. The number of hub taxa was significantly correlated with temperature (R^2^ = 0.975, p = 0.005), with a high of 71 hubs in the +9 °C network (Figure 3D). Archaeal hubs were present in the +4.5 °C, +6.75 °C and +9 °C networks, and fungal hubs were observed in all networks except the +2.25 °C network (Figure 3D). Class-level taxonomic distribution of the hub taxa revealed that *Acidobacteriae* were the most prominent across all treatments and depths, followed by the *Verrucomicrobiae* (Figure 4). In the +9 °C network, six unique classes were represented in the hubs, including the methanogenic class *Methanosarcinia* (Figure 4). Two unique classes were present in the +0 °C network hubs, *Myxococcia* and *Dehalococcoidia* (Figure 4). Only three of the fungal hubs could be assigned to a guild by FUNguild analysis, including possible ectomycorrhizal and saprotrophic *Agaricomycetes* hubs and a *Sordariomycetes* hub that is a probable plant pathogen.

**Figure 4:**
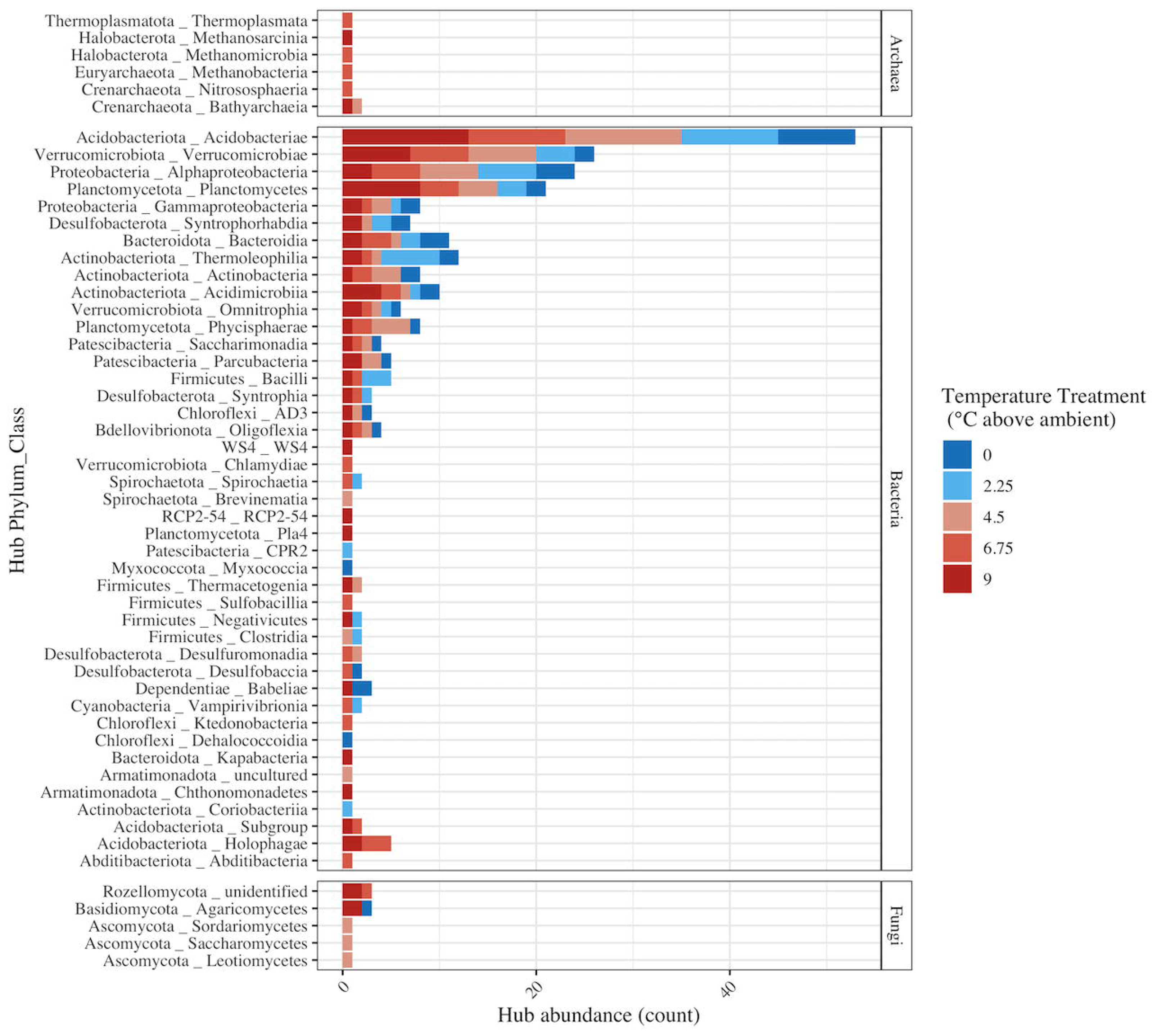
Class-level distribution of network hubs across temperature treatments (color). Hubs were aggregated at class-level taxonomic annotations (y-axis) and faceted by kingdom-level taxonomy.

### 3.4 Peat soil mass loss and compositional changes are highly variable

On average, the mass of peat soil in decomposition ladders was significantly lower after three years of treatment (x ± SD mass loss = 4.50% ± 11.07%; Kruskal-Wallis χ^2^ = 6.7158, p = 0.0095) (Figure 5A); however, no significant effects of depth, temperature, CO_2_ treatment, or their interactions on peat soil mass loss were observed (Figure 6A; Table S1). Peat C and N content decreased significantly over the course of the experiment (Figure S7), and C:N of the peat was significantly higher at T_f_ compared to T_0_ (Kruskal-Wallis χ^2^ = 7.0841, p = 0.0078) (Figure 5B), indicating a more rapid loss of N compared to C. Similar to soil mass loss, neither depth nor the treatment variables had a significant effect on the change in C:N (Figure 6B; Table S1).

**Figure 5:**
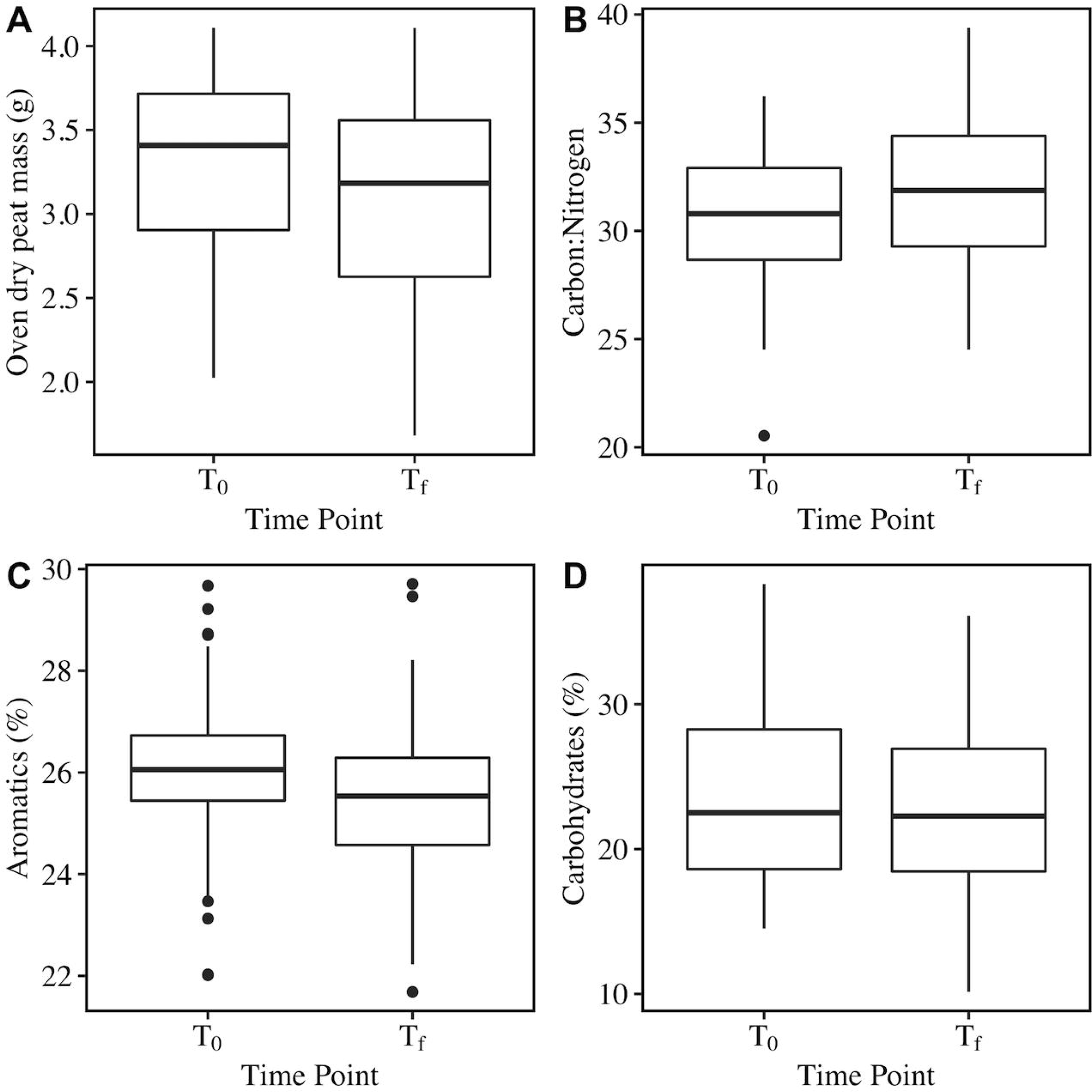
Comparisons of peat soil oven dry mass (A), carbon:nitrogen (B), percent aromatics (C), and percent carbohydrates (D) in peat decomposition ladders at the start of the experiment (T_0_) and after three years of incubation in the SPRUCE enclosures (T_f_). Box and whisker plots display the median (middle of box), quartiles (top and bottom of box), minimum and maximum values excluding outliers (end of whisker) and outliers (points).

**Figure 6:**
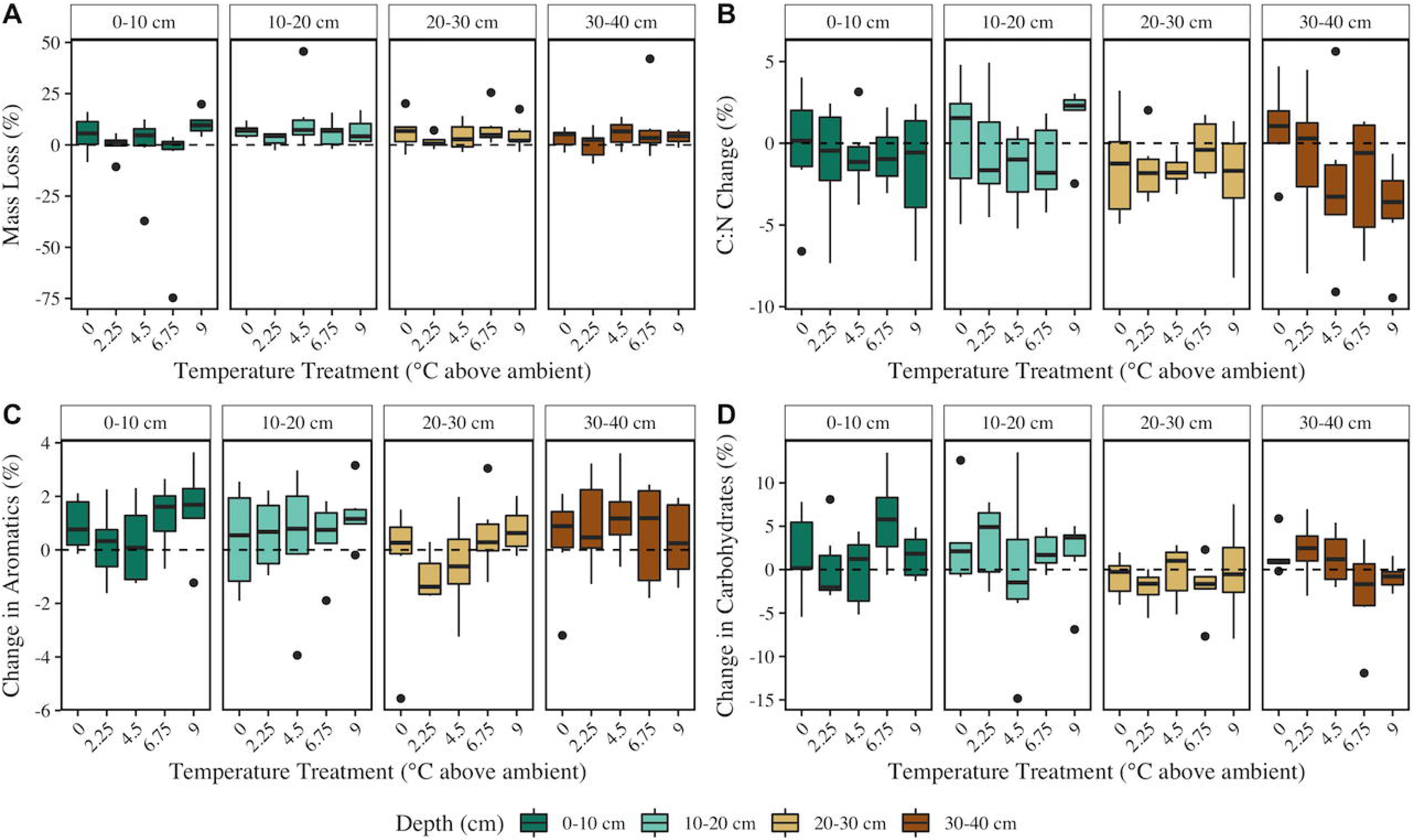
Peat mass loss (% of starting mass lost) (A), percent C:N change (T_f_ – T_0_) (B), percent aromatics change (T_f_ – T_0_) (C), and percent carbohydrates change (T_f_ – T_0_) (D) across depths (facets) and temperature treatments (x-axis).

Results of Fourier-transform infrared spectroscopy (FTIR) analyses further showed limited effects of the treatments on peat soil composition after three years of incubation. Aromatics (%) decreased significantly over the course of the experiment (Figure 5C), but there was no significant effect of soil depth or treatment on the change in aromatics (Figure 6C; Table S1). Depth significantly impacted the change in % carbohydrates, with largest change in the % carbohydrates at 0-10 cm. However, there was no significant difference in % carbohydrates between T_0_ and T_f_ (Figure 5D), and temperature and CO_2_ treatment did not significantly impact carbohydrate content (Figure 6D; Table S1).

## 4. Discussion

The effects that climate change will have on peatland microbial communities and the resulting impacts on soil decomposition have yet to be fully resolved but could have outsized effects due to the massive stores of carbon in peatlands. Previous results from SPRUCE have shown rapid loss of carbon presumably driven by increased decomposition in response to elevated temperature [10]. In this study, we utilized decomposition ladders to investigate the effects of warming and elevated CO_2_ on peat microbial communities and decomposition while limiting inputs from primary productivity. Our results show that bacterial/archaeal and fungal communities are significantly impacted by the SPRUCE treatments; however, in contrast to previous research, we did not observe a significant effect of temperature, elevated CO_2_, or soil depth on mass loss or compositional changes. An average of less than 4.5% of the initial mass was lost over the three-year experiment regardless of treatment, demonstrating the high recalcitrance of organic soils in peatlands, and suggesting that these prior reported results may driven by turnover of more recently fixed C rather than the historic C stocks of peat studied here. These small differences also illustrate just how inherently difficult measure such decomposition is to measure given unavoidable experimental variability between replicates and other sources of error, combined with small mass losses of these recalcitrant substrates.

Bacterial/archaeal and fungal community compositions were significantly influenced by the SPRUCE treatments (Figure 1, Table 1), indicating that increased temperature and atmospheric CO_2_ as a result of climate change may alter microbial ecology in peatlands. Consistent with previous research, bacterial/archaeal community responses to temperature treatment were more pronounced than fungal responses [22, 48, 49]. Bacterial/archaeal α-diversity was significantly highest in the warmest SPRUCE enclosures across all soil depths, whereas the influence of temperature treatment on fungal α-diversity was depth specific and only observed at q=2 (Figure 2, Table 2). Ecosystem disturbance and environmental stress have been negatively correlated with microbial diversity [50], thus higher bacterial/archaeal diversity in warmed enclosures suggests that warming may alleviate environmental stress on prokaryotic communities in peatlands. Higher bacterial/archaeal diversity may be a direct cause of warming; however, contrasting results have also been observed in anaerobic peat soil microcosms that investigated direct warming effects [17, 21]. Indirect effects such as warming-induced increases in substrate and nutrient availability [51] are therefore more likely driving changes in bacterial/archaeal diversity by partially relieving nutrient stress. Our experimental design did not allow us to delineate between direct and indirect effects of warming, as porewater total organic carbon and nutrient concentrations were largely correlated with temperature treatment on average over the course of the experiment (Table S2). However, previous research has demonstrated that labile substrate and nutrient availability and microbial diversity are positively correlated which supports our results.

Network analyses further revealed the impact of increased temperature on peat microbial community structure. The number of nodes and edges were highest in the networks from warmed enclosures (Figure 3), and a positive correlation between the number of input ASVs and edge and node counts indicates that this is likely a reflection of species richness. Species richness and changes in network topology such as a decreased ratio of positive to negative edges and increased modularity in microbial networks have been shown to correspond to higher community stability [50, 52, 53], potentially through increased microbial functional redundancy [54]. Paired with diversity measures, our network results indicate that warming may positively influence microbial community stability.

The abundance of microbial hub taxa (those taxa which are highly connected within the network) was positively correlated with temperature treatment (Figure 3D). The number of hub taxa within microbial networks has been associated with functional potential of the microbial community [55, 56], and hub taxa may exert strong influence over microbiome structure and ecosystem function regardless of their relative abundance within the community [57, 58]. Therefore, our results suggest that warming may promote increased microbial functional potential in these peatland ecosystems.

We observed *Methanomicrobia*, *Methanobacteria*, and *Methanosarcinia* hubs in the +6.75 °C and +9 °C networks (Figure 4), supporting results that show methanogenesis is an increasingly important function in peatland carbon-cycle response to warming. The relative abundance of methanogenic taxa has been shown to increase in response to increasing temperature in incubations [17], and rates of methane production have largely been shown to increase with warming [59], including in incubation studies of soil from the S1 bog [17].

Detection of an acetoclastic methanogen in the +9 °C network corresponds to isotopic analysis of CH_4_ in the SPRUCE enclosures indicating that acetoclastic methanogenesis is increasing with warming [59]. A significant linear relationship between *in situ* porewater concentrations of CH_4_ and temperature treatment has been observed in the top 25 cm of soil at SPRUCE [12], further supporting our results. Methane has a global warming potential of 28 times that of CO_2_ on a 100- year time span [6], and higher potential for methanogenesis in response to climate change may fuel a positive feedback loop.

Methanogenesis can be supported by syntrophic interactions, especially under nutrient limiting conditions that are observed in bogs [60, 61]. Two known syntrophic taxa, *Syntrophia* [62] and *Syntrophorhabdia* [63], were identified as hubs within the networks (Figure 4), suggesting the potential importance of syntrophy within the sites. Other hubs identified in the networks from warmed enclosures including *Bathyarchaeia* and *Holophagae* may further support increased acetoclastic methanogenesis, as these taxa have been previously shown to have the potential for acetogenesis in anoxic environments [64, 65].

Studies of peatland responses to disturbance often investigate fungal and bacterial/archaeal communities independent of one-another despite knowledge of the complex interplay across microbial domains [66, 67]. Here, we observed the highest number of fungal nodes and hub taxa in networks from decomposition ladders in warmed enclosures, suggesting that peatland warming may promote trans-domain interactions, further arguing for such holistic approaches. Possible saprotrophic and ectomycorrhizal fungal hub taxa were primarily observed in the networks from warmed enclosures apart from an *Agaricomycetes* hub in the +0 °C network (Figure 4). Fungal communities play an important role in organic-matter (OM) decomposition in peatlands [68], and warming may favor dominance of saprotrophic and mycorrhizal fungi from *Basidiomycota* and *Ascomycota* [69], as partially observed in our study. Additionally, warming treatments in these systems are inextricably linked to drying of the peat surface and increased depths to water table. It is thus possible that peat drying and water table changes, more so than warming, may be responsible for the shifts in fungal communities observed; however, the average water table height was similar across all SPRUCE enclosures in the week prior to termination of the experiment (0.17 m ± 2.47 cm) as well as the three prior months.

In contrast to bacterial/archaeal diversity, fungal α-diversity was significantly lower under elevated CO_2_ compared to ambient. Fungal responses to CO_2_ treatment are likely mediated by plant responses, as CO_2_ treatment at SPRUCE is above-ground and is unlikely to directly alter soil biogeochemistry. Interactions between plant roots and fungi are common, and fungi are most prevalent (absolute abundance) near the surface of the peatland where active plant growth occurs [70]. Our results are intriguing and suggest further investigation into plant-fungal interactions under elevated CO_2_ conditions.

Significant changes in the peat microbial communities in response to SPRUCE treatments were not mirrored by the peat soil decomposition rates. We anticipated that increased temperature would result in increased peat soil mass loss, as temperature treatment at SPRUCE has resulted in rapid carbon loss that was presumed to be driven by enhanced decomposition [10], increased CO_2_ and CH_4_ in porewaters [12], and increased microbial respiration of solid phase peat [13]. However, our results showed that soil mass loss and C:N were not significantly impacted by temperature or CO_2_ treatment over the course of three years (Figure 6). The lack of differences is likely driven by the short time scale and low initial mass of peat soil in the decomposition bag study compared to the SPRUCE enclosures. Hanson et al. (2020) estimated the rate of carbon loss from SPRUCE to be 31.3 g Cꞏm^−2^ꞏyear^−1^ꞏ °C^−1^using approaches including elevation changes and ecosystem CO_2_ flux mass balances. Using this rate to estimate the expected loss of carbon from the peat decomposition ladders suggests that differences in mass loss across temperature treatments were on the order of milligrams over a three-year period, thus likely requiring a level of precision that we were unable to be obtain in our litter bag-based experiments.

On average, only 4.5% of the initial peat soil mass was lost over the course of the experiment (Figure 5). The low mass loss demonstrated the recalcitrance of the organic soils in the decomposition ladders is likely driven by a combination of factors including the anoxic, acidic, and oligotrophic conditions of the sites, as well as the chemical composition of the peat. Peat soils at SPRUCE are largely derived from *Sphagnum*, which is known to engineer acidic, nutrient poor, waterlogged conditions [71, 72] and produces anti-microbial compounds and metabolites [73, 74], thereby inhibiting microbial degradation processes. Even the mass loss of fresh *Sphagnum* litter in decomposition bags has been previously shown to be similarly low with only ∼10% of initial mass lost after two years [75, 76], so these lower rates for peat soil should not be unexpected.

Diffusion of exogenous dissolved organic matter (DOM) into the decomposition ladders may have also help explain the differences in results between mass loss and community change. Dissolved organic matter is preferentially mineralized by peatland microorganisms when compared to solid-phase peat [77], and fresh plant inputs of DOM can even fuel microbial respiration in deep peat soils [78]. Utilization of exogenous DOM may also partially explain discrepancies between our results and previous results from SPRUCE that have shown increased CO_2_ and CH_4_ production with warming. Porewater concentrations of total organic carbon were highest in the + 9 °C enclosures near the termination of our experiment, and higher inputs of DOM may have led to increased OM mineralization and shifts in microbial community structure without impacting peat soil mass in the decomposition ladders.

Similar to soil mass-loss observations, FTIR analysis indicated that temperature and CO_2_ treatment had no effect on changes in peat soil composition. The relatively short duration of the experiment may have masked temperature effects on the percent aromatics and carbohydrates of the peat, although previous results have indicated that carbon at SPRUCE is compositionally stable [79]. We are hopeful that our future planned ladder extractions at our site with their longer field incubation periods may allow for better assessment of treatment effects on peat soil decomposition.

## Conclusions

While we did not observe changes in peat soil mass or composition across the SPRUCE treatments in this study, much previous research from the SPRUCE experiment has shown loss of OM near the surface and increased greenhouse gas production in response to elevated temperatures. Our results suggest that these losses in OM may be driven by changes in microbial community structure and dynamics, as we observed significant changes in microbial diversity and network structure in response to warming. The apparent decoupling of changes in peat soil mass and composition and microbial communities may be limited by the very slow peat decomposition rates and precision of mass loss estimates in our study. Collectively, our results and previous results from the SPRUCE experiment therefore suggest that climate change may alter peatland microbial ecology however the ultimate effects of these changes in on rates of degradation of OM and greenhouse gas production in boreal regions remains unclear at this time.

## Acknowledgements

We would like to thank Randy Hedin and Evelyn Magner (US Forest Service Northern Research Station) for their assistance in assembling and testing the peat decomposition ladders. This research was sponsored by the Environmental Systems Science Program, U.S. Department of Energy, Office of Science, Biological, and Environmental Research, as part of the Terrestrial Ecosystem Science Scientific Focus Area at Oak Ridge National Laboratory. Oak Ridge National Laboratory is managed by UT-Battelle, LLC, for the U. S. Department of Energy under contract DE-AC05-00OR22725. This research used resources of the Compute and Data Environment for Science (CADES) at the Oak Ridge National Laboratory, which is supported by the Office of Science of the U.S. Department of Energy under Contract No. DE-AC05- 00OR22725.

## Author contribution statement

Conceptualization – NAG, PJH, RKK, CWS; Investigation – SR, RKK, NAG, KCO, AAC, AS, JPC, DMK; Formal analysis – SR; Original draft writing – SR; Draft revision – SR, RKK, NAG, KCO, AAC, AS, JPC, DMK, CWS

## ORCID

SR: 0000-0001-9559-1154

NAG: 0000-0003-0068-7714

RKK: 0000-0002-6419-8218

PJH: 0000-0001-7293-3561

KCO: 0000-0002-0875-2720

AAC: 0000-0003-1142-4709

DMK: 0000-0002-4307-2560

AS: 0000-0001-8269-9270

JPC: 0000-0002-3303-9708

CWS: 0000-0001-8759-2448

