## Supplemental figures and tables for "Elevated temperature alters microbial communities, but not decomposition rates, during three years of in-situ peat decomposition"

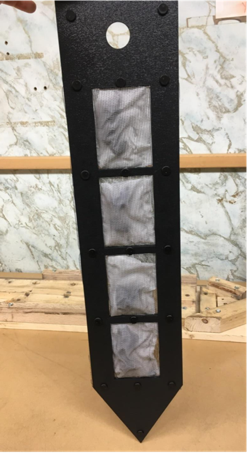

Figure S1: Peat decomposition ladder prior to deployment into the SPRUCE enclosures.

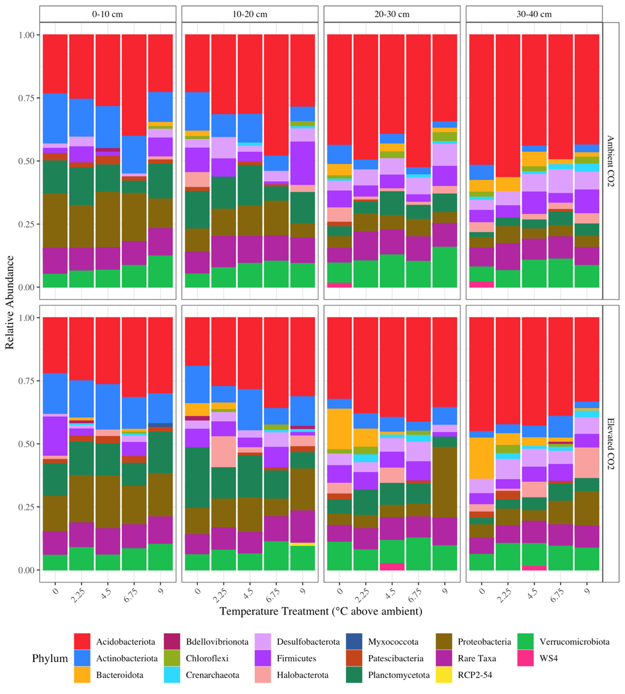

Figure S2: Phylum-level taxonomic distribution of prokaryotic taxa in decomposition ladders across depths (horizontal facets), CO_2_ treatments (vertical facets), and temperature treatments (x-axis). Bars represent the mean relative abundance of each phylum across replicate samples within each depth X temperature treatment X CO_2_ treatment.

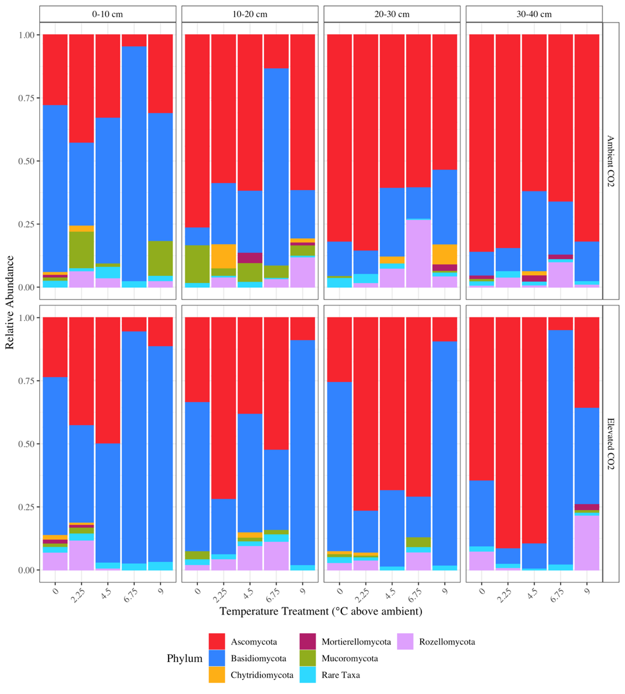

Figure S3: Phylum-level taxonomic distribution of fungal taxa in decomposition ladders across depths (horizontal facets), CO_2_ treatments (vertical facets), and temperature treatments (x-axis). Bars represent the mean relative abundance of each phylum across replicate samples within each depth X temperature treatment X CO_2_ treatment.

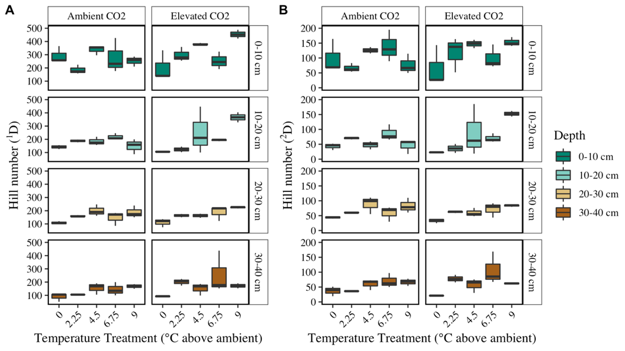

Figure S4: Bacterial/archaeal α-diversity at q = 1 (Shannon diversity) (A), and q = 2 (inverse Simpson diversity) (B) across depths (vertical facets/colors), CO_2_ treatments (horizontal facets), and temperature treatments (x-axis).

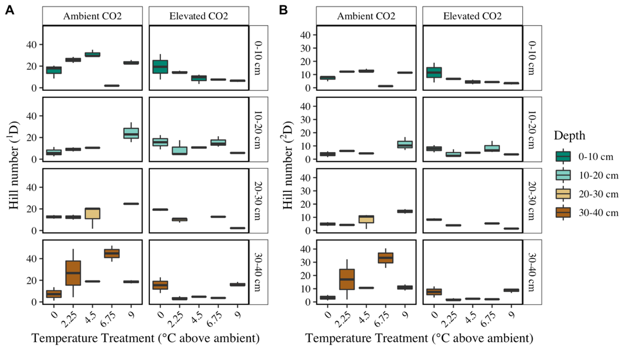

Figure S5: Fungal α-diversity at q = 1 (Shannon diversity) (A), and q = 2 (inverse Simpson diversity) (B) across depths (vertical facets/colors), CO_2_ treatments (horizontal facets), and temperature treatments (x-axis).

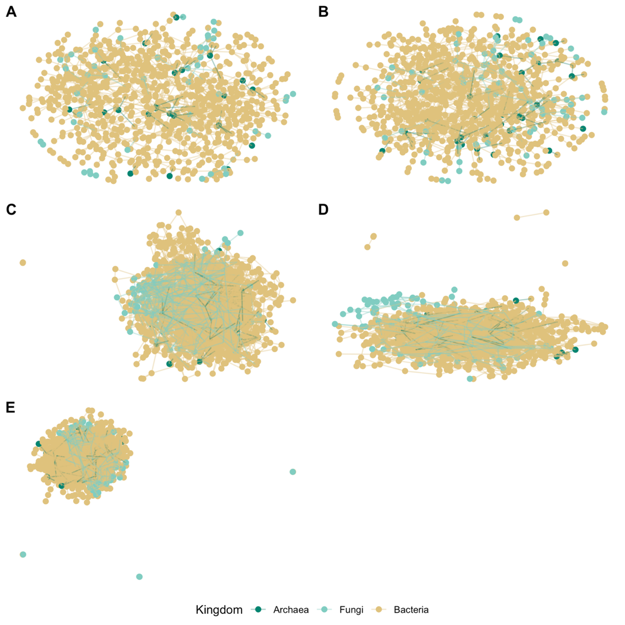

Figure S6: Trans-domain microbial networks constructed from +0 ºC plots (A), +2.25 ºC plots (B), +4.5 ºC plots (C), +6.75 ºC plots (D), and +9 ºC plots (E). Nodes (points) are colored by based on kingdom level taxonomic assignments. Edges are represented by lines connecting nodes. All replicate samples across all depths were included in each temperature treatment network (n = 24 samples/network). All networks were visualized using the Fruchterman-Reingold layout method.

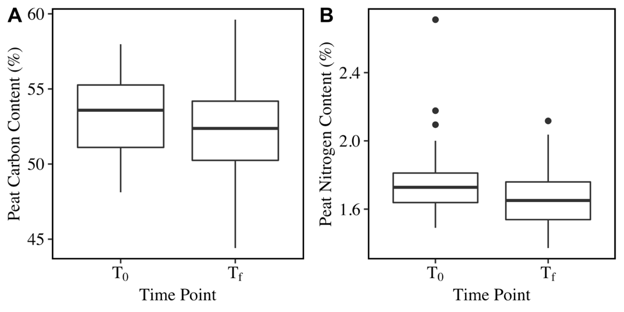

Figure S7: Comparisons of peat carbon content (%) (A) and peat nitrogen content (%) (B) in peat decomposition ladders at the start of the experiment (T_0_) and after three years of incubation in the SPRUCE enclosures (T_f_). Box and whisker plots display the median (middle of box), quartiles (top and bottom of box), minimum and maximum values (end of whisker) and outliers (points).

Table S1. P values associated with linear models were used to assess the effect of depth, temperature treatment, and CO2 treatment on peat soil mass loss and chemical composition changes. Only the effects of depth upon carbohhydrates was significant (P<0.05)

|  | **Mass loss** | **C:N change** | **Aromatics change** | **Carbohydrates change** |
| --- | --- | --- | --- | --- |
| **Depth** | 0.17 | 0.27 | 0.07 | **0.02** |
| **Temperature** | 0.66 | 0.14 | 0.06 | 0.57 |
| **CO2** | 0.5 | 0.63 | 0.32 | 0.11 |
| **Depth*Temperature** | 0.97 | 0.11 | 0.5 | 0.22 |

Table S2. Porewater geochemical profiles at three depths within each SPRUCE enclosure (0 cm, 20 cm, 50 cm). Data are averaged across duplicate temperature treatment enclosures and across multiple sampling events during the course of the experiment (n sampling events = 33). Raw data are accessible at Griffiths, N. A., S.D. Sebestyen, K.C. Oleheiser, J.M. Stelling, C.E. Pierce, E.A. Nater, B.M. Toner, & R.K. Kolka. 2016. SPRUCE Porewater Chemistry Data for Experimental Plots, Beginning in 2013. Oak Ridge National Laboratory, TES SFA, US Department of Energy, Oak Ridge, Tennessee, USA. Version 3. <https://doi.org/10.3334/CDIAC/spruce.028>.

| **Depth** | **Temp** | **PH** | **Specific conductivity (µS/cm)** | **Chloride (mg/L)** | **Sulfate (mg/L)** | **Calcium (mg/L)** | **Potassium (mg/L)** | **Magnesium (mg/L)** | **Sodium (mg/L)** | **Total Aluminum (mg/L)** | **Total Iron (mg/L)** | **Total Manganese (mg/L)** | **Silicon (mg/L)** | **Strontium (mg/L)** | **Ammonium N (mg N/L)** | **Nitrate N (mg N/L)** | **Soluble reactive phosphorus (mg P/L)** | **Total Nitrogen (mg N/L)** | **Total phosphorus (mg P/L)** | **Total organic carbon (mg C/L)** |
| --- | --- | --- | --- | --- | --- | --- | --- | --- | --- | --- | --- | --- | --- | --- | --- | --- | --- | --- | --- | --- |
| **0 cm** | **0** | 3.55 | 54.11 | 0.40 | 0.23 | 1.68 | 0.54 | 0.58 | 0.99 | 0.35 | 1.65 | 0.02 | 3.00 | 0.01 | 0.20 | 0.00 | 0.07 | 1.34 | 0.11 | 65.35 |
|  | **2.25** | 3.51 | 69.22 | 0.40 | 0.31 | 2.78 | 1.03 | 0.90 | 1.09 | 0.64 | 4.53 | 0.06 | 3.27 | 0.01 | 0.35 | 0.01 | 0.11 | 1.60 | 0.14 | 105.26 |
|  | **4.5** | 3.41 | 94.84 | 0.82 | 0.42 | 2.99 | 1.92 | 1.33 | 1.65 | 0.86 | 4.57 | 0.05 | 5.01 | 0.02 | 0.25 | 0.01 | 0.07 | 1.98 | 0.12 | 137.63 |
|  | **6.75** | 3.48 | 88.24 | 0.85 | 0.41 | 2.83 | 3.70 | 1.10 | 1.37 | 0.82 | 4.18 | 0.09 | 4.88 | 0.01 | 1.01 | 0.02 | 0.48 | 2.76 | 0.58 | 131.58 |
|  | **9** | 3.43 | 95.89 | 0.74 | 0.56 | 3.07 | 3.60 | 1.15 | 1.65 | 0.83 | 3.68 | 0.03 | 5.18 | 0.02 | 0.59 | 0.02 | 0.22 | 2.38 | 0.29 | 137.61 |
| **20 cm** | **0** | 3.55 | 48.83 | 0.13 | 0.19 | 2.10 | 0.12 | 0.66 | 1.23 | 0.51 | 1.15 | 0.02 | 7.22 | 0.01 | 0.02 | 0.00 | 0.01 | 1.02 | 0.04 | 63.18 |
|  | **2.25** | 3.49 | 63.16 | 0.30 | 0.20 | 2.99 | 0.33 | 0.99 | 1.32 | 0.66 | 1.61 | 0.03 | 9.62 | 0.01 | 0.23 | 0.01 | 0.09 | 1.34 | 0.11 | 93.55 |
|  | **4.5** | 3.42 | 69.80 | 0.43 | 0.31 | 2.82 | 1.01 | 0.94 | 1.49 | 0.79 | 2.02 | 0.02 | 9.10 | 0.01 | 0.24 | 0.01 | 0.09 | 1.55 | 0.13 | 98.70 |
|  | **6.75** | 3.47 | 81.38 | 0.76 | 0.26 | 2.62 | 1.69 | 0.83 | 1.34 | 0.84 | 1.56 | 0.02 | 11.99 | 0.01 | 2.02 | 0.01 | 0.87 | 3.47 | 1.03 | 123.43 |
|  | **9** | 3.49 | 77.01 | 0.64 | 0.36 | 3.90 | 1.58 | 1.17 | 1.51 | 0.88 | 2.24 | 0.03 | 12.50 | 0.02 | 0.56 | 0.01 | 0.15 | 1.96 | 0.19 | 129.59 |
| **50 cm** | **0** | 3.59 | 48.62 | 0.15 | 0.14 | 2.38 | 0.36 | 0.69 | 1.32 | 0.60 | 1.00 | 0.01 | 7.94 | 0.01 | 0.01 | 0.00 | 0.01 | 1.03 | 0.03 | 64.82 |
|  | **2.25** | 3.54 | 54.02 | 0.22 | 0.14 | 2.46 | 0.40 | 0.77 | 1.36 | 0.58 | 0.88 | 0.02 | 11.86 | 0.01 | 0.34 | 0.00 | 0.02 | 1.54 | 0.05 | 73.27 |
|  | **4.5** | 3.59 | 49.50 | 0.25 | 0.11 | 2.34 | 0.86 | 0.64 | 1.33 | 0.59 | 1.05 | 0.01 | 10.79 | 0.01 | 0.40 | 0.00 | 0.01 | 1.41 | 0.03 | 63.70 |
|  | **6.75** | 3.56 | 65.89 | 0.53 | 0.17 | 2.26 | 0.89 | 0.64 | 1.37 | 0.84 | 0.99 | 0.01 | 13.29 | 0.01 | 1.96 | 0.01 | 0.49 | 3.52 | 0.56 | 99.65 |
|  | **9** | 3.54 | 64.02 | 0.42 | 0.22 | 2.87 | 0.89 | 0.85 | 1.52 | 0.84 | 1.18 | 0.02 | 15.07 | 0.02 | 1.00 | 0.01 | 0.16 | 2.46 | 0.19 | 100.04 |
